# Resilience assessment in complex natural systems

**DOI:** 10.1101/2023.09.12.557305

**Authors:** Camilla Sguotti, Paraskevas Vasilakopoulos, Evangelos Tzanatos, Romain Frelat

## Abstract

Ecological resilience is the capability of an ecosystem to maintain the same structure and function and to avoid crossing catastrophic tipping points. While fundamental for management, concrete ways to estimate and interpret resilience in real ecosystems are still lacking. Here, we develop an empirical approach to estimate resilience based on the stochastic *cusp* model derived from catastrophe theory. Our *Cusp Resilience Assessment* (CUSPRA) has three characteristics: i) it provides estimates on how likely a system is to cross a tipping point characterized by hysteresis, ii) it assesses resilience in relation to multiple external drivers, and iii) it produces straightforward results for ecosystem-based management. We validated our approach using simulated data and demonstrated its application using empirical time-series of an Atlantic cod population and of marine ecosystems in the North and the Mediterranean Sea. We show that CUSPRA provides a powerful method to empirically estimate resilience in support of a sustainable management of our constantly adapting ecosystems under global climate change.

## Introduction

Ecological resilience, the capability of an ecosystem to maintain the same structure and function under the influence of external drivers, and hence to avoid crossing catastrophic tipping points, is one of the emerging concepts of sustainability^1,2^. While maintaining ecosystems or populations resilient to global cumulative stressors is fundamental for ecosystem health, we are lacking concrete and simple ways to empirically estimate and interpret resilience in real ecosystems^3–5^. This shortcoming is partly due to inconsistency related to the resilience concept (existence of different definitions across disciplines), but especially due to the complexity of the natural systems^6–8^. The lack of solid methodologies to empirically estimate resilience has caused its neglect in ecosystem-based management frameworks. For instance, European maritime policies (such as the European Union’s Marine Strategy Framework Directive-MSFD) have not embraced the concept, even though the understanding of a system’s resilience would facilitate the adoption of more adequate management measures^9–12^.

Resilience is usually discussed within the concept of regime shifts, i.e. non-linear and abrupt transitions of a system between alternate states that differ in configuration and/or properties^4,13,14^. Indeed, a highly resilient system should be able to withstand more pressures, staying clear from tipping points and associated regime shifts^14,15^. As a result, systems with low resilience need to be managed with caution in order to avoid transitions to unwanted configurations^16^. By contrast, if a system is locked within an undesirable state (e.g. an overfished ecosystem), actions to erode its resilience would be needed to facilitate a shift towards a desirable state^4^. Knowledge about past resilience dynamics and regime shifts of ecosystems would help policy makers understand whether a system has undergone abrupt regime changes and to identify the external drivers^17–20^. Importantly, reliable empirical resilience estimates are crucial for assessing the ability of a system to recover to previous baselines, often a key objective of ecosystem-based management, or for detecting if the new state is potentially irreversible^21,22^.

While measuring resilience is relatively straightforward from experiments or theoretical models, the challenge lies in determining resilience from empirical data for large ecological systems that cannot be experimentally manipulated^8,23–25^. A number of methods have been proposed to estimate resilience in aquatic populations and ecosystems^8^. Early warning signals of regime shifts estimate resilience based on mathematical properties of time series representing the systems^3,26,27^. Other approaches to estimate resilience in ecosystems are based on species interactions and make use of properties of network analyses and food-web modelling^28–30^. Lastly, another approach is to quantify the resilience of alternate states in probabilistic terms using information from a large number of different systems^31,32^ or simulated system states^33^. A common issue across all these methods is that they require large and detailed (empirical or simulated) datasets with high temporal resolution, which are usually unavailable for complex natural systems.

An empirical approach for the detection of regime shifts and quantification of resilience that makes use of typical ecological monitoring time series is represented by the Integrated Resilience Assessment (IRA)^34,35^. IRA quantifies resilience by identifying two or more multiple stable states of a system, affected by one driver that acts as a system stressor, and measuring how far the system is from both the steady state and a tipping point^34–38^. This method has been applied across different marine ecosystems^34,35,37,39,40^. IRA has however the shortcoming that the underlying statistical model is not representing discontinuous regime shift dynamics including hysteresis and potential irreversibility^34^. Moreover, IRA is a single driver approach and hence a strong simplification of complex ecological systems that are usually affected by multiple and possibly interacting cumulative drivers^41^. Models based on catastrophe theory such as the stochastic cusp model (*cusp)* can better represent such discontinuous dynamics^8,42–44^. *Cusp* is a non-linear modelling approach based on a cubic differential equation which allows to identify tipping points derived from a bifurcation in a system depending on two interactive drivers, and to test for reversibility of system states^43–46^. Implementing *cusp* into IRA would hence be a step forward in developing a reliable resilience assessment from empirical data.

Here, we combined the *cusp* and concepts from IRA to develop a *Cusp Resilience Assessment* (*CUSPRA*) that has three characteristics: i) it provides estimates on how likely a system is to cross a tipping point characterized by hysteresis, ii) resilience is assessed in relation to multiple external drivers, and iii) it produces straightforward results for ecosystem-based management. These three characteristics are fundamental for the development of a method that is both easy to apply on typical empirical datasets and that facilitates a better understanding on how systems respond to external drivers. We validate *CUSPRA* using artificial data and demonstrate its application using empirical time-series data related to an Atlantic cod population and ecosystem dynamics in the North Sea and the Mediterranean Sea. We show that CUSPRA provides an advanced methodology to empirically estimate resilience in support of a sustainable management of our constantly changing and adapting ecosystems under global climate change.

## Material and Methods

### Building the resilience indicator

We built *CUSPRA* combining the stochastic *cusp* model with principles derived from the resilience quantification approach of IRA^34,43^. CUSPRA uses a 4-step approach: (1) select the variables to use in the model, (2) fit and evaluate the *cusp* model, (3) estimate resilience, and (4) assess the results in relation to the interaction of two drivers.

**Step 1: fitting the stochastic cusp model**

To estimate resilience in populations and ecosystems we firstly need to fit the stochastic cusp model to the data. The stochastic cusp model was developed by the mathematician Rene Thom in the 1980s and allows to model the dynamics of a state variable depending on two interactive drivers^8,42–44,47^. The model is based on a cubic differential equation (Eq.1) extended with a Wiener’s process to add stochasticity (Eq.2)^43,45^.

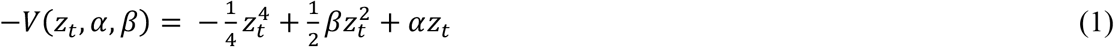

where *V*(*z*_*t*_, *α, β*) is a potential function whose slope represents the rate of change of the system (the system state variable is called z_t_), depending on the two control variables (α,β)^43^.

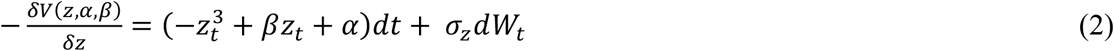

The two control variables, α and β, are the factors that can cause the rise of non-linear discontinuous dynamics. α is the asymmetry variable that controls the dimension of the state variable (the variable we want to model such as a time series of an ecosystem or a population). Therefore, if α increases the state variable will change accordingly. Instead, β is the so-called bifurcation variable and can modify the relationship between z_t_ and α from linear and continuous to non-linear and discontinuous^45^. In the model α, β and z are modelled as linear function of the original variables we want to fit in the model such as fishing, temperature (SST), population state (Eq.3)^43,45^. A combination of different drivers can also be used to model the three factors fitted in the stochastic cusp model.

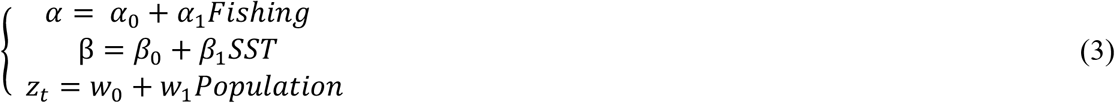

where α_0_, β_0_ and w_0_ are the intercepts, and α_1_, β_1_ and w_1_ the slopes of the models.

The stochastic cusp model is thus able to detect three different types of dynamics of the state variable, linear and continuous, non-linear and continuous and discontinuous^45^. To do that a Cardan’s discriminant is calculated (Eq.4).

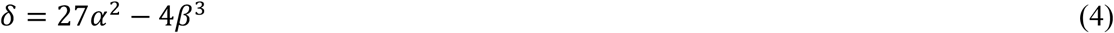

If the Cardan discriminant is smaller or equal to zero, then the state variable follows non-linear discontinuous dynamics and has multiple equilibria, while if it is higher than zero the system will follow continuous dynamics^48^. The model results can be plotted in a 2D plot showing the type of dynamics of the state variable depending on the level of the two control factors (SI Fig.1)^45^. The Cardan’s discriminant is represented in this 2D plot and indicates where the linear dynamics occur and where the combination of the two drivers creates the bifurcation, and thus the non-linear discontinuous dynamic emerge. In the bifurcation area, called cusp area (i.e. the area below the fold), three equilibria are possible, two stable, and one unstable. This is the transition area where the tipping point is present. Therefore, the model can determine the combination of the two drivers that lead to discontinuous dynamics.

The stochastic cusp model directly fits two alternate models, one linear (Eq.5) and one logistic (Eq.6) in order to select between different dynamics of the system.

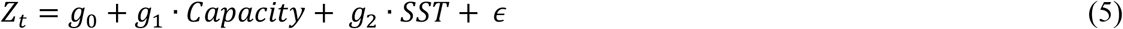

Where g_0_ is the intercept of the model, and g_1_ and g_2_ the slopes coefficients of the two control variables and *ϵ* is the normally distributed random error (mean=0, variance= σ square).

The logistic model, showing non-linear but continuous dynamics was fitted as:

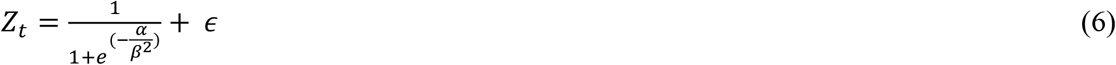

Where Z, α and β are canonical variables of the observed state and control variables defined in Eq.3 and *ϵ* is the zero mean random disturbance.

Before proceeding with the resilience estimation, it is necessary to evaluate the model and check that the stochastic cusp model is the better model compared to the two alternative ones. The model evaluation can be done using the pseudo R-squared. However, the simple comparisons between R squared could mask the better fit of the cusp model^48^. Therefore, to confirm that the best model is the cusp three other indicators need to be checked: (1) The percentage of points inside the cusp area needs to be more than 10%; (2) The state variable needs to have a multimodal nature thus needs to present a change point; (3) the estimate of the state variable needs to be significant. If these criteria are met, the stochastic cusp model is superior to all the other models and is a good simplification of the state variable dynamics and thus it is possible to proceed with the resilience estimation^45,48^.

**Step 2: Estimating resilience, building from the IRA**.

Resilience is estimated as for the IRA, by calculating the distance of the current state to the instability area using two components, a vertical and a horizontal component^34^. The resilience is high when the observed control variables (α, β) are far from the bifurcation area while the resilience is low when they (α, β) are within the bifurcation set. By definition from Eq.4, the bifurcation set is defined by (Eq. 7):

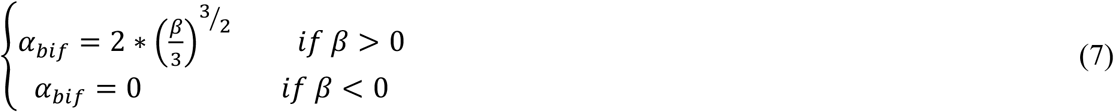

The horizontal component of resilience is the distance of *α* from the bifurcation set or instability area (Eq.8).

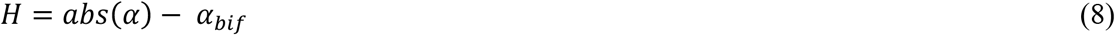

H is negative when inside the bifurcation set (abs(α)< *α*_*bif*_, low resilience) and positive when outside the bifurcation set (higher resilience).

The vertical component is the distance to linearity, defined by the bifurcation variable *β*, representing how discontinuous is the system (Eq.9).

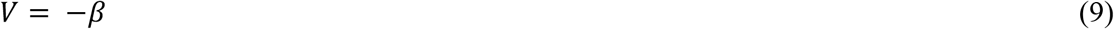

V is negative when discontinuous and positive when linear (β<0).

The resilience (Ri) is then the weighted average between the horizontal and the vertical component. We give double weight to the horizontal component (H) to further stress the importance of *α* for the resilience of the system in comparison to the vertical component (Eq.10).

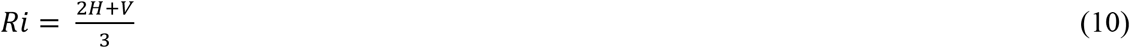

Highly negative Ri indicates a highly discontinuous system in the bifurcation set, while highly positive values indicate a linear system far from the bifurcation set. We transformed this resilience value (Ri) using hyperbolic tangent transformation to get an indicator of resilience (RA) between 0 and 1, with 0 reflecting low resilience and 1 high resilience (Eq. 11).

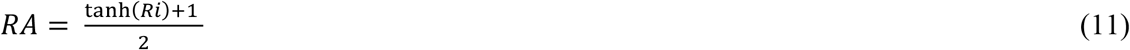

In this way, we obtained an indicator that can be comparable across multiple systems and multiple populations. The resilience estimation is computed for every point of the state variable, depending on the two control variables. Thus, if we fit as a state variable the time series of the biomass of a population, for every point in time of it, a value of RA will be calculated. A low *RA* (value close to 0) means that with regard to the fluctuation levels of the stressors the system is unstable, i.e. large changes in state can happen with little changes in the stressor variables and that the system is close to a tipping point. On the contrary, a large *RA* (close to 1) indicates that the system is mostly linear and thus if the control variables change the system will change linearly (i.e. the system is far from the tipping point). Being based on the cusp and the IRA, our concept of resilience does not test for resilience per se (as a “condition” of the system independently of the variables that may affect it) but estimate the resilience depending on two external drivers and determine how close a system lies to a tipping point.

**Step 3: testing the indicator**

To test whether the indicator provides meaningful results, we run some simulations. We created state variables from multiple stochastic cusp models having the α variable being strongly autocorrelated (AR1=0.9) and the β variable being normally distributed. The mean of β varied in five different model scenarios, from -2 (mostly linear) to 2 (mostly discontinuous) with an increment of 1. We ran ten repetitions of each scenario with a time series of 50 time-steps.

We tested three hypotheses to validate our resilience metric:

H1) the resilience decreases with the level of non-linearity (β)

H2) a state with low resilience has higher probability of large changes

H3) a state with low resilience is linked to anomalously large variance

H1 helped us to show how the model works, while H2 and H3 validate that our concept of resilience corresponds to the one described in the literature, where a low resilient system exhibits higher variance and higher probability to change.

To test these hypotheses, we estimated the CUSPRA resilience on each simulation. For H1, we compared the minimum RA to the set level of non-linearity (β). We expected that the minimum RA decreases with non-linearity so it should increase with the mean level of β. For H2, we compared the absolute difference between successive state values with the value of RA. We expected that absolute differences would decrease with higher RA. For H3, we computed the variance within a 5-time step windows, and we compared it to the level of RA. We expected that the variance is low for high resilience, and that the variance increases when resilience decreases.

### Data

To test the newly developed indicator we used data from four published scientific articles using the stochastic cusp model or the IRA and ranging from populations to community and to traits configurations^35,36,45^. We built a Shiny App to allow all members of the scientific community to test our resilience indicator with their own data (https://rfrelat.shinyapps.io/CUSPRA/). In all the four examples the drivers tested to assess resilience were fishing and climate change (i.e. Sea Surface Temperature), derived from different sources. The examples are thus comparable. To test population resilience, we used data from the Atlantic cod (*Gadus morhua*) stock of the the North-East Arctic Sea, from Sguotti et al., 2019 ^45^. We use as a proxy for the state of Atlantic cod in the North-East Arctic, Spawning Stock Biomass (SSB) data deriving from stock assessment. Fishing mortality (F), also estimated from stock assessment, and Sea Surface Temperature (SST) were used as drivers to test the resilience (for more details see Sguotti et al., 2019). The data ranged from 1966 to 2016^45^ (SI Fig.2).

Ecosystem resilience was explored using North Sea and Mediterranean Sea data. In this case, to build a proxy of the community firstly a multivariate reduction technique (PCA) needed to be applied^35^. CUSPRA, similarly to both IRA and the stochastic cusp model, needs to fit a single vector (a time series) representing the system state to the model. The North Sea community index was built performing a PCA to a matrix of data coming from the Continuous Plankton Recorder (CPR) and International Bottom Trawl Survey (IBTS) ^49^. SST derived from NOAA ErSST v5 and fishing effort from Couce at al. 2019, were used as drivers to test resilience^68,69^. The data ranged from 1985 to 2019 (for more info Sguotti et al., 2022a) (SI Fig.3) ^49^. In the Mediterranean case example, a community dataset was built using landings data extracted from the database of the Food and Agriculture Organization (FAO) of the United Nations and then reduced using a PCA^35^. As drivers, fishing capacity estimated for the entire Mediterranean Sea by FAO and SST from the NASA PODAAC were used. The data ranged from 1985 until 2013 (SI Fig.4). Finally, to estimate resilience of the traits of the Mediterranean Sea community, we used landings traits data derived from Tsimara et al. 2021. This dataset was derived by combining the Mediterranean fisheries landings data of the FAO for the years 1985–2015 (http://www.fao.org/fishery/statistics/GFCM-capture-production/en) with a matrix including data on 23 complete traits related to the biology, ecology, trophic role, distribution, habitat and behaviour of 205 species (mostly fish, but also molluscs and crustaceans) found in Koutsidi et al. (2019)^70^. The multiplication of the two datasets resulted into a matrix of community weighted mean traits landings by year in the Mediterranean Sea that was analysed in Tsimara et al. (2021) (SI Fig.5).

All the analyses were performed in R (version 4.0.2) using the packages cusp^48^ for the cusp modeling. The implementation of the calculation of the resilience indicator (RA) is documented in the github repository (https://github.com/rfrelat/Cuspra). A Shiny app was created and made available to help scientists calculate the resilience indicator on their time series or run other simulations (https://rfrelat.shinyapps.io/CUSPRA/).

## Results

### Evaluating system dynamics

We built *CUSPRA* following a 4-step approach: (1) select the variables to use in the model, (2) fit and evaluate the *cusp* model, (3) estimate resilience, and (4) assess the results in relation to the interaction of two drivers (Fig.1). The first two steps deliver a *cusp* model as the basis for the resilience estimation. The *cusp* represents discontinuous changes in a dynamic system depending on two control drivers (SI Fig.1)^45,47^. The system (i.e. a state variable) is represented by a vector of data, typically a time-series, and can be anything from abundance estimates of a population to proxies of ecosystem state(s), depending on the system whose resilience we are interested to analyze (Fig.1). The dynamics of the state variable depending on two interactive drivers are modelled with a cubic differential equation^42,43,48^. An asymmetry factor α controls the state of the system, thus inducing system changes, for instance from a regime A to a regime B. This factor is usually a linear model of an external driver that can be managed such as fishing effort in harvested ecosystem systems, or a combination of external drivers^45,49,50^ (Fig.1). A second bifurcation factor β modifies the relationship between the asymmetry factor and the system state from linear continuous to non-linear and discontinuous, and thus shapes the functional form of system dynamics, i.e. if the system approaches the bifurcation. The bifurcation factor is usually a linear model of an external driver that induces changes in the system, such as temperature, eutrophication, or climate indices or a combination of all those^45,49–51^ (Fig.1).

**Figure 1.**
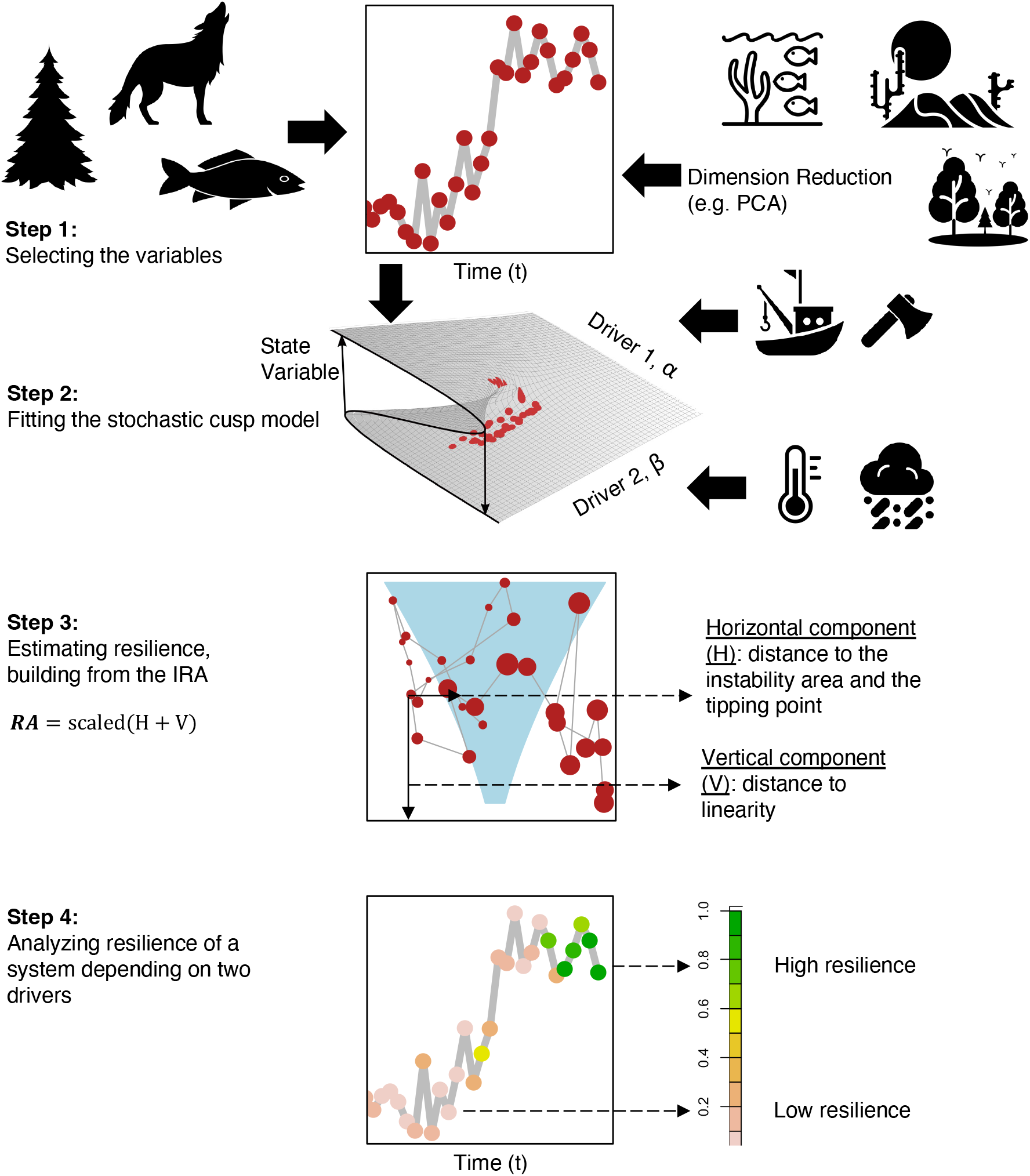
The 4-steps of CUSPRA. *1) Selecting the variables:* The state variable should be a time series, that can represent anything from a single population to an ecosystem. If multivariate time-series are available, these need to be reduced to an ecosystem metric using a dimension reduction technique such as Principal Component Analysis (PCA). *2) Fitting the stochastic cusp model:* The model, using a cubic differential equation, represents the dynamics of a system from linear and continuous to non-linear and discontinuous, depending on the combination of the two drivers. *3) Estimating resilience (RA)*: Projecting the results of the cusp model in 2D, an estimate of resilience can be calculated for every state in time depending on the two drivers. Resilience is calculated based on the vertical distance of a point to linearity and on the horizontal distance of a point to the cusp area (in light blue) or tipping point (see also Fig. 3). In the cusp area three equilibria, two stable and one unstable, are possible, in the area below the fold. *4) Analyzing resilience depending on the two interactive drivers:* CUSPRA allows to understand how the resilience changes over time and to compare the results between case study systems.

The solution of the cubic differential equation of the model can either detect the multiple equilibria in the state variable, indicating discontinuous dynamics and a regime shift, or identify that just one equilibrium exists, and thus linear continuous dynamics prevail (see Methods)^43,45,46,50^. The model is generally represented in a horizontal plain determined by the combination of the two drivers (Fig.1, step 2,3). The system can move in the plain from areas in which just one equilibrium exists, i.e. linear dynamics, to areas with multiple equilibria, i.e. discontinuous dynamics (i.e. a regime shift, the light blue area) (SI Fig.1). This area is also identified as the area below the fold, where tipping points can occur and resilience is low (SI Fig.1)^43,48^. Once the cusp model has been fitted and an extensive model validation procedure reveals the data to represent true discontinuous dynamics including tipping points and hysteresis (see methods), the resilience estimation can be performed (Fig.1, Step 3).

### Resilience estimation

Step 3 in *CUSPRA* entails the resilience estimation based on the 2D representation of the *cusp* (SI Fig.1). Resilience is assessed based on principles of the IRA, that measures it as the sum of distances from the stable state and tipping point^34,35^. In *CUSPRA*, resilience is also calculated based on two components. First, the vertical distance (V), which is the distance between the state variable and the part of the plain representing linear dynamics (Fig.1 step 3, Fig.2a). V is determined by changes in the bifurcation variable (β), the factor controlling the type of the system dynamics. The second component is the horizontal distance (H), the distance between the state of the system and the instability/cusp area, i.e. the area where the three states are possible (Fig.1 step 3, Fig.2b). Resilience inside this transition area is supposed to be really low since it corresponds to the area below the fold. These two quantities were summed in an equation and then transformed using hyperbolic tangent transformation to obtain an overall resilience estimate (RA) scaled between 0 and 1 (Fig.2c,d, see Methods). The empirical estimate (RA) provided by step 3 eventually allows step 4 in *CUSPRA*, the analysis on how likely the system is to switch state and to cross a tipping point. If RA is close to 0, resilience is low, and the system is in a highly transitory state and thus close to a tipping point. Instead, if RA is close to 1, resilience is high, and the system is in a stable state and far away from a tipping point (Fig.2).

**Figure 2.**
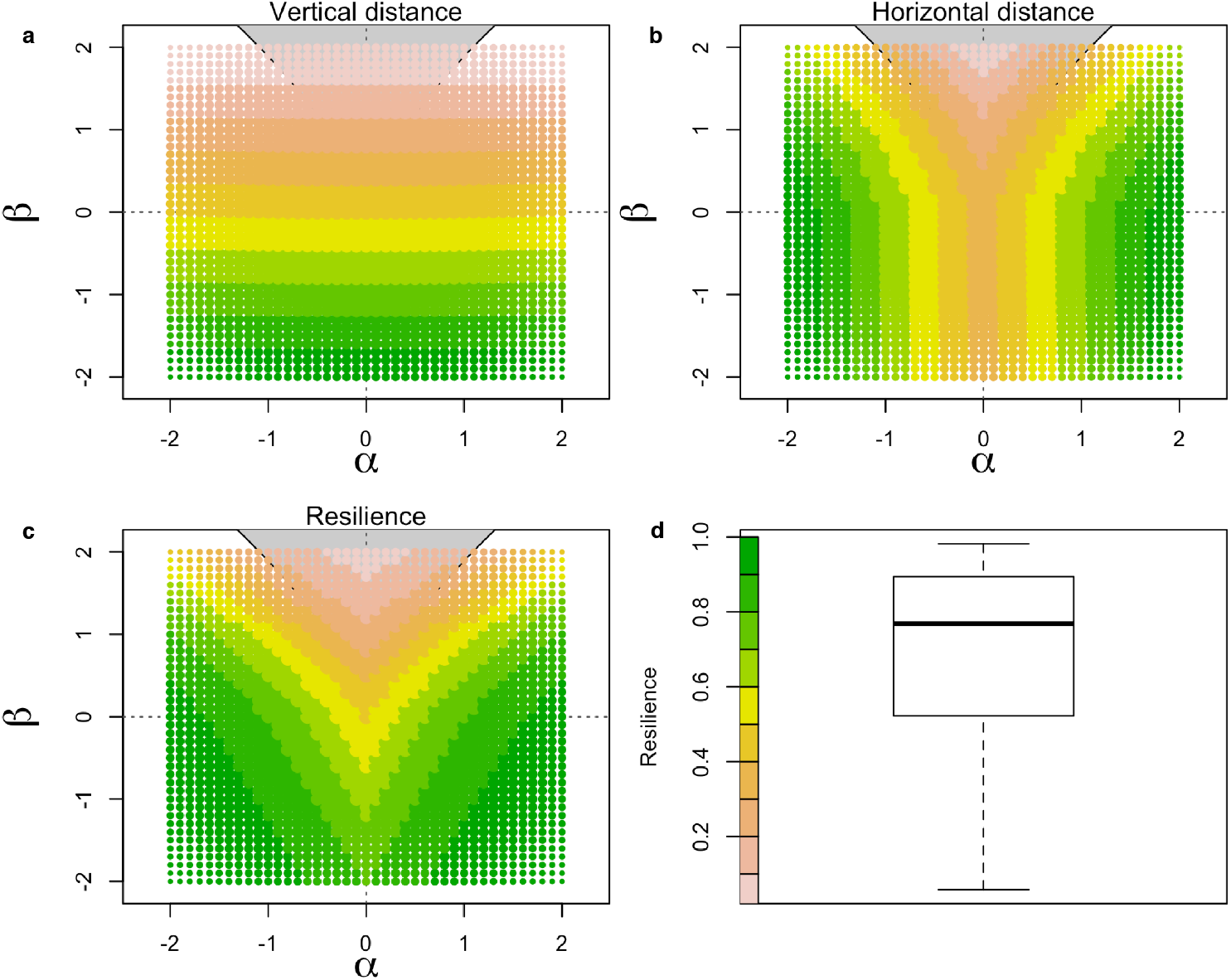
The CUSPRA resilience indicator (RA) | The 2D representation of the stochastic cusp model; for a-c: on the x axis, α, the driver controlling the dimension of the state variable; on the y axis, β, the driver controlling whether the relationship between α and the state variable is linear and continuous or non-linear and discontinuous. In grey the instability area determined by the combination of α and β values where three equilibria are possible. The colors of the areas of the 2D panel, correspond to different resilience estimates (pink= 0 and green =1). **a**) The vertical component of resilience where resilience is estimated as the distance from linearity. **b**) The horizontal component of resilience estimated as the horizontal distance from the cusp area. **c**) The CUSPRA resilience indicator obtained by summing the vertical and horizontal resilience estimates. The resilience is lower inside the instability area. **d**) The CUSPRA resilience values and a boxplot showing the RA distribution in the 2D area.

### Validation

We validated our CUSPRA method by simulating time series of two drivers, α and β, to derive a state variable from the stochastic cusp model. We assumed the α variable to be strongly autocorrelated (AR1=0.9) and the β variable to be normally distributed, to imitate, respectively, an anthropogenic driver such as fishing, and a climate variable such as temperature or productivity. The mean of β varied in five different model scenarios, from -2 (mostly linear) to 2 (mostly discontinuous). We ran 10 repetitions of each scenario with a time series of 50 time-steps. With these simulations we tested three hypotheses to evaluate whether our model was meaningful. Firstly, we wanted to show that resilience decreases with the level of non-linearity (β). A β of – 2 has high resilience while a β of 2 has a resilience close to 0 indicating that the choice of a meaningful bifurcation variable to estimate resilience in real examples is crucial (Fig. 3a). The second and the third hypotheses were related to the concept of resilience itself, i.e. a system with low RA has a higher probability of change and is linked to larger variances. These two properties are related to the concept of critical slowing down, a property typical of systems approaching a tipping point^27,52^. Using our simulated data, we confirmed that the amount of change of the state variable between two time-points t and t_+1_ (delta Δ) increases and becomes more variable as RA declines (Fig. 3b). Moreover, at a low RA the variance of the state variable (calculated with a 5-year moving window) increased (Fig. 3c). These results indicate that the resilience estimated by CUSPRA corresponds to the concept of resilience in ecology^3,8,53,54^.

**Figure 3:**
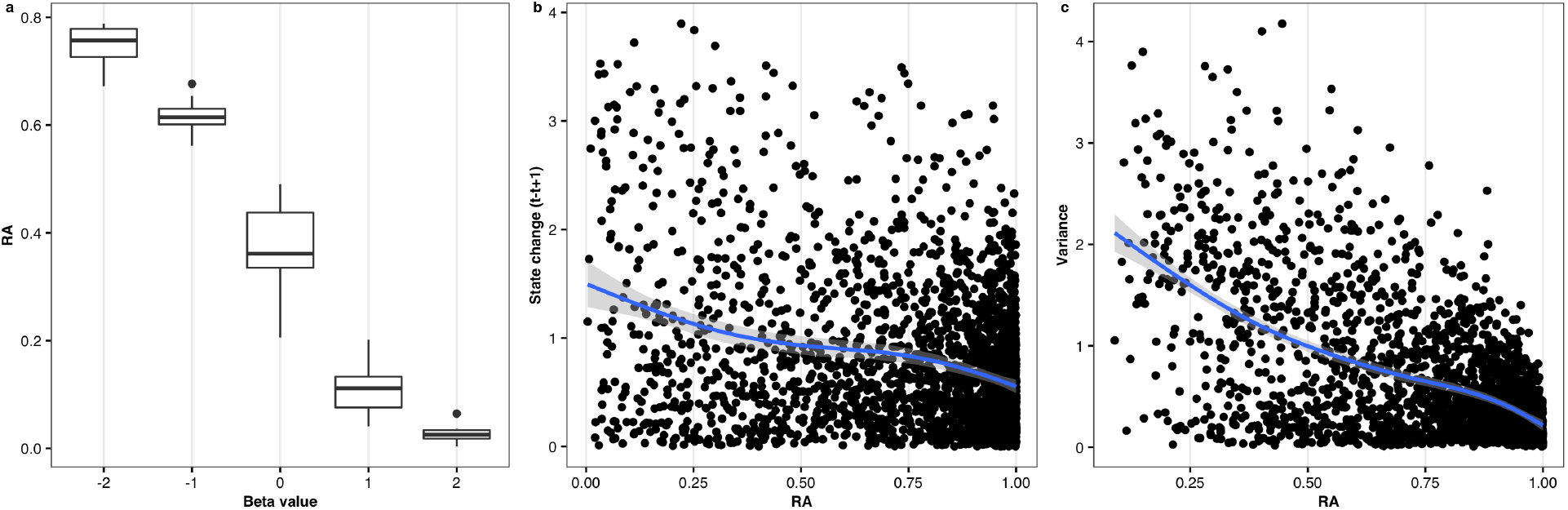
Simulation results for model evaluation. a) Distribution of the minimum RA resilience (y-axis) for simulated time series with different mean values of the β drivers (x-axis). **b**) Changes of the state variable depending on its resilience (x-axis). The blue line represents the smoothed relationship between these two variables, showing that at lower RA values the probability of change increases. **c**) Variance of the state variable depending on its resilience value RA calculated with a 5-year moving window. The blue line represents the smoothed relationship showing that variance decreases with increasing resilience.

### Application to real systems

We used four examples to illustrate the application of our new resilience estimation approach. First, we estimated resilience of a commercially important fish stock (Atlantic cod, *Gadus morhua*, in the North-East Arctic^45^) based on fish stock assessment data (modelled population outputs). We used Spawning Stock Biomass (SSB; the weight of mature fish in the stock) as the predictor for the state variable, while α was predicted by fishing mortality (F), and β by Sea Surface Temperature (SST) (for more details about the modelling see Sguotti *et al*., 2019, SI Fig.2). The cusp model fitted well and was superior to the alternative linear model (SI Tab.1). While the logistic model was superior based on the R squared, all the other criteria (e.g. points inside the bifurcation area, etc) were met and thus the cusp model fitted well to the data. The analysis shows that at the beginning of the time series the resilience of the stock was low (Fig.4a,b). During this period the stock biomass was low, fishing mortality on a medium level and temperature low as well. The subsequent increase in fishing pressure pushed the stock towards a resilient and hence stable (albeit undesirable) low biomass state, outside the cusp area (Fig.4a, b green, small dots). Decreasing fishing mortalities after 2007 coupled with an increase in temperature eroded the resilience of the low biomass state (pink dots), moved the system back into the cusp area, and caused a tipping point towards a high biomass state c. in 2009. A further increase of temperature subsequently increased the resilience of the high biomass state (Fig.4a, b). The North-East Arctic cod provides an example in which a management measure (i.e. reduced fishing pressure) eroded the resilience of the unfavorable low biomass state, and in combination with increasing temperatures moved the stock towards a more favorable stable state, representing a positive transition.

Next, we applied CUSPRA to the dynamics of entire ecosystems (North Sea and Eastern Mediterranean Sea) represented by multivariate data sets. To estimate resilience in these cases, we initially needed to reduce the dimensionality of the data, here through a Principal Component Analysis (PCA; see Sguotti *et al*., 2022 and Vasilakopoulos *et al*., 2017, SI Fig.3,4) and use the first mode of variability derived from the PCA (PC1) to predict the state variables in cusp models. We predicted α with time series of fishing pressure (see Methods) and β with SST. The cusp models fitted well and were superior to the alternative models based on the R squared and the additional screening were passed (SI Tab.1). The analyses revealed very low resilience in the North Sea ecosystem at the beginning of the time series (Fig. 4c,d) when fishing pressure was high and temperature low, and the ecosystem in a state dominated by cod^49^. With decreasing fishing pressure (mainly in the 2000s) and increasing temperature (mainly in the 2010s) the North Sea ecosystem changed from low to high resilience due to structural changes in terms of species composition^49^. On the contrary, the Eastern Mediterranean Sea ecosystem shifted from a high into a low resilience state, also reflecting a change in community structure, mainly due to increased fishing pressure (Fig.4e,f). Recently, increasing temperatures are maintaining low resilience in the Eastern Mediterranean Sea, while the structure of the fish community changed (big dots)^35^.

**Figure 4:**
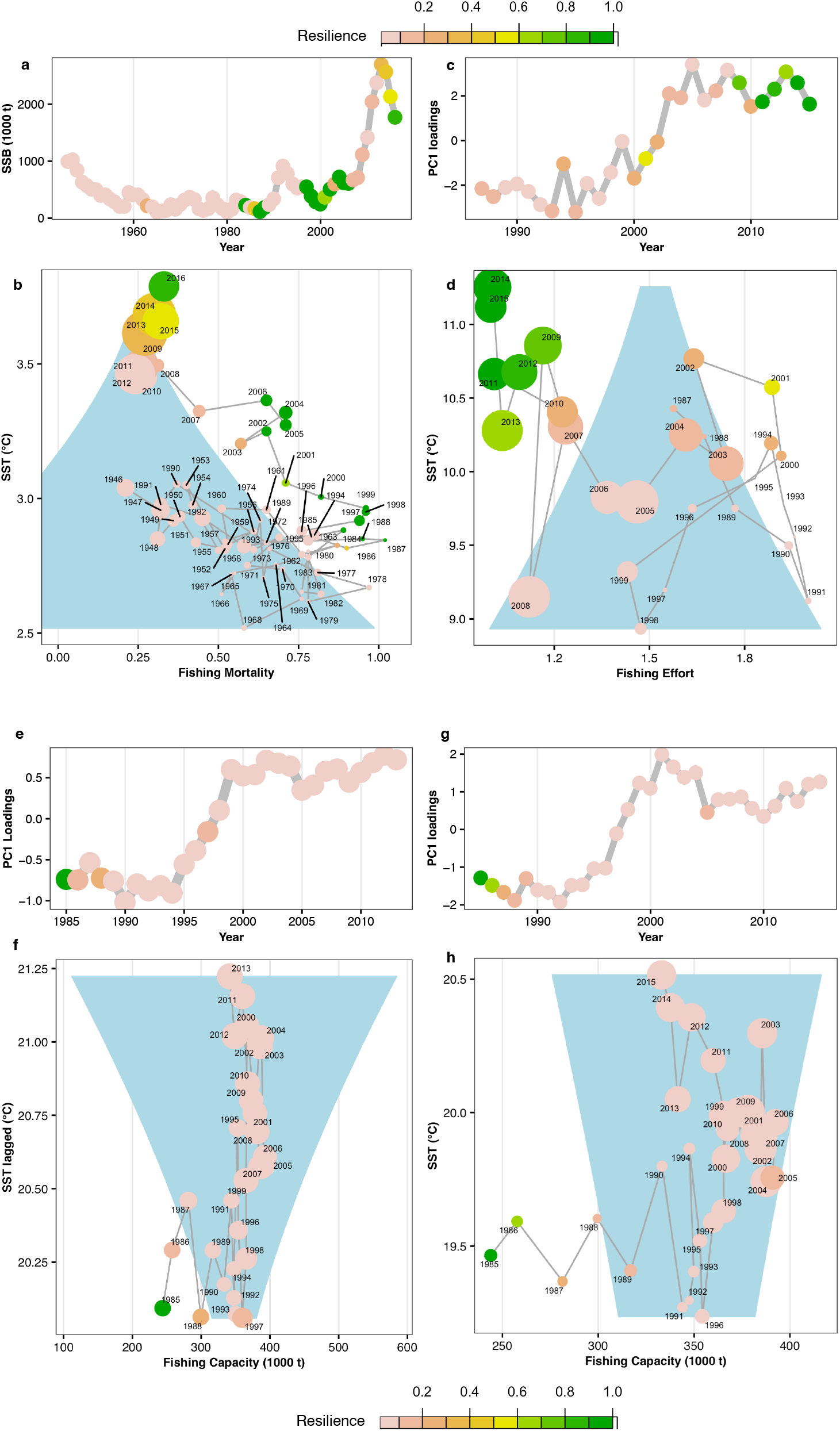
The CUSPRA application. a,b) Resilience of North-East Arctic cod spawning stock biomass (SSB) depending on fishing mortality and Sea Surface Temperature (SST). (c,d) Resilience of the North Sea ecosystem depending on fishing effort and SST. (e,f) Resilience of the Eastern Mediterranean ecosystem depending on fishing capacity and SST (lagged by 2 years). (g,h) Resilience of the Mediterranean ecosystem based on traits depending on fishing capacity and SST. A,c,e,g, show the time series of resilience in the different systems. b,d,f,h show the resilience values in the 2D representation of the cusp model with the blue area corresponding to the cusp area where 3 equilibria are possible, i.e. two stable and one unstable. The dimension of the dots is proportional to the magnitude of the state variable. The color of the dots corresponds to the color of our CUSPRA resilience estimate RA, pink= 0, low resilience, green= 1, high resilience.

Finally, we used CUSPRA to estimate resilience of the trait composition of the fish community of the Mediterranean Sea^36^. PC1 here represents the main mode of variability of the trait space of the fish community and was used to predict the state variable in the cusp model. Again, we used a measure of fishing pressure and SST to predict the α and β variables, respectively (Si Fig.5). Again, the cusp model fitted well and was superior to the alternative models (SI Tab.1). Similar to the analysis of the Eastern Mediterranean fish community, resilience decreased initially with increasing fishing pressure (Fig.4g,h). With the decline of fishing capacity and especially the increase in temperature, the system abruptly shifted to a new state (see also Tsimara *et al*. 2021) in the late 1990s, being however of low resilience, i.e. prone to further shifts. The analysis denotes that only a drastic reduction of fishing pressure would be able to drive the system towards a resilient state, and the increase of SST will require more drastic management approaches.

## Discussion

Here we have developed CUSPRA, a new method to assess resilience of ecological systems based on empirical data. Our method allows to estimate how close a system is to a tipping point and hence to a shift into a new state or regime. Our objective was to build a method that estimates resilience to tipping points, i.e. shifts between regime changes characterized by hysteresis or even irreversibility. Such dynamics are represented in CUSPRA through the application of the stochastic cusp model^55^, a mathematical approach that can detect bifurcation in a system and thus identify tipping points^8,44,45,50^. Moreover, CUSPRA estimates resilience based on the effect of multiple interacting drivers and provides an indicator directly applicable in ecosystem-based management. We validated our approach using simulated data and tested our newly developed resilience indicator using four empirical example data sets comprising three different system types, i.e. a fish population, two fish communities and a trait configuration^35,36,45^. Interestingly, by using the same drivers in all case studies, we can understand their varying impacts in the different systems. For instance, the increase in temperature in the North Sea ecosystem, coupled with strong management measures (i.e. decreased fishing pressure)^49,56^ has led to a new resilient ecosystem state, while in the Mediterranean Sea similar changes in temperature but an increasing exploitation caused a new state but with low resilience. In the Mediterranean case, fishing pressure appears to be still excessively high and the system is likely closer to its maximum thermal tolerance ^57,58^. Our examples demonstrate that CUSPRA is useful to understand the resilience of a system and how close it is to tipping points. Our new method not only advances empirical studies in resilience science but can also be directly applicable in ecosystem management settings, even beyond the marine environment, since it is easily transferable to a variety of systems, not only ecological systems but also socio-ecological, financial or behavioral systems.

Another characteristic of our approach is that resilience is quantified based on the interaction of multiple stressors and thus allows to quantify hysteresis to interactive pressures^44,45^. This is a step forward compared to other methods that estimate resilience without accounting for system drivers, or methods such as the IRA, that estimate resilience depending on only one driver^27,34^. In the Anthropocene, multiple drivers acting in an interacting or cumulative fashion are increasingly likely to impact our resources and ecosystems, so methods that can consider a higher level of complexity are more suitable to model real systems^59,60^. It is possible to fit in the CUSPRA even more than two drivers adding variables in the two factors controlling the system states. These would allow for a multidimensional study of resilience^51^. Finally, the estimation of a resilience indicator ranging from 0 to 1 allows the CUSPRA to be easily translatable into management and to be used for comparisons across multiple systems. The simplicity of the final indicator is appealing for management purposes since it constitutes a straightforward, unitless metric indicating whether the system under management is in a state that is rather stable or prone to change. Moreover, the possibility of comparing and understanding the resilience of different populations or ecosystems to the same stressors, or even of completely different systems, can improve our knowledge about resilience and favor a better comprehension of this complex concept. Thus, our new method will translate the multifaceted concept of resilience in an easily comprehensible metric that can be used in many different disciplines providing useful information for management purposes. Examples on how to translate the concept into management can be found in SI Tab.2.

Our CUSPRA approach takes the quantitative resilience assessment of complex natural systems based on empirical data a step forward. While our new method better represents resilience based on the effect of two interactive drivers, it also shows some limitations linked with the stochastic cusp model regarding the consideration of autocorrelation in the time series of the variables and concerning model evaluation^45,48^. Several indicators hence need to be evaluated before a cusp model can be assessed to be superior to alternative linear and continuous models^45,48^, i.e. before CUSPRA can be used to assess resilience. In all our four examples the cusp model was found to provide more reliable fits to the data than alternative models, which indicates that these ecological systems exhibited non-linear and discontinuous dynamics.

The model also presents opportunities to be improved. At present, as most of the empirical models, our approach is “data-driven” and thus it is difficult to make predictions. Indeed, the method can be used to estimate resilience of a system in hindsight only, while it cannot provide inferences about its future development^45,50^. Nevertheless, inspecting the interactive effects of drivers and the position of the state variable relative to the instability area provides information on the likely development in the near future. Other future developments could be to extend CUSPRA to the possibility of detecting more than two stable states. This “limitation” resides in the cusp model that is only able to depict two alternative states and hence no multiple consecutive tipping points can be assessed. Other bifurcation models, such as the butterfly should be able to detect more than two stable states and thus could also be integrated in CUSPRA^44^. Nevertheless, usually empirical time series are short, and it is reasonable to believe that only two states of the system likely exist in the data time scales. If longer time series are to be fitted to the data, an extensive data analysis before fitting the CUSPRA is necessary. Visualizing the development of the state variable will show if fitting two separate CUSPRA models is necessary^61^. CUSPRA models very complex phenomena in a simplistic way^8^. While the strong simplification might be criticized, it is one of the strengths of this approach. The simplicity makes it easily translatable into management. Nonetheless, a CUSPRA analysis should always be supported by additional indicators to be able to better understand the system state.

### How could CUSPRA be used in ecosystem-based management settings?

First of all, knowledge on the type of dynamics that the system has shown in the past (linear or discontinuous) is important to understand if a system is vulnerable to regime shifts^9^. Knowing which drivers have caused the discontinuous dynamics, can give an indication on the resilience of the system towards these drivers and thus how vulnerable the system is to them^4,62^. This is important for management since it can help to establish which drivers need to be managed in a more precautionary approach in order to enhance the resilience of the system and avoid the crossing of a tipping point^63^. Moreover, CUSPRA can determine the levels of the stressors at which the system will switch into a new state, i.e. the position of the tipping point. This is an important knowledge for managers since it can help to mitigate stressors in order to avoid their critical levels^4,64^. Finally, the method gives a snapshot of the resilience of the system at present and thus can indicate whether a system will approach a tipping point in the immediate future. This is a particularly important information in order to decide whether to adopt precautionary approaches or whether to try to restore the system towards initial conditions ^4^. Potentially, building CUSPRA with multiple drivers and their combinations could help managers to define a safe operating space of the system and thus favor better management approaches^65,66^. Estimating resilience is fundamental to properly manage natural systems, however this concept is seldomly included in management due to methodological limitations^1,2^. CUSPRA, by estimating resilience of ecological systems impacted by multiple stressors, allows for a better quantification of resilience and a direct application to management which is urgently needed if we want to manage constantly changing and adapting systems under global climate change^22,67^.

## Acknowledgements

CS was funded by the EU HORIZON RESET (Resilience Estimation to SET management goals in marine ecosystems) Project (101065994) under the HORIZON-MSCA-2021-PF and by the BMBF-funded project SeaUseTip, Spatio-temporal analysis of tipping points in the socio-ecological system of the North Sea (funding code: 01LC1825A-C). This work is also a contribution to the EU H2020 COMFORT project that received funding from the European Union’s Horizon 2020 research and innovation program under grant agreement number 820989 (project COMFORT, Our common future ocean in the Earth system–quantifying coupled cycles of carbon, oxygen, and nutrients for determining and achieving safe operating spaces with respect to tipping points). The work reflects only the author’s/authors’ view; the European Commission and their executive agency are not responsible for any use that may be made of the information the work contains. We acknowledge the ICES Working Group COMEDA (Comparative Ecosystem-based Analyses of Atlantic and Mediterranean marine systems) under which this study was started and developed. We would like to thank Prof. Christian Moellmann for his precious advices and his brilliant suggestions while developing the study.

## Author’s contribution

CS and RF conceived the study. CS and RF carried out all the analyses and produce the figures. CS, RF, PV, ET discussed the methods and the approaches. All authors helped drafting the manuscript and gave comments and inputs.

## Conflicts of interests

There were no conflicts of interest.

## Data and materials availability

All data and codes are stored in the GitHub repository (https://github.com/rfrelat/Cuspra) and are freely accessible. A Shiny App was also developed to allow other researchers or stakeholder to easily try the method with their data or simulated data (https://rfrelat.shinyapps.io/CUSPRA)

## Notes

### Competing Interest Statement

The authors have declared no competing interest.

