## Supplementary Information for "Resilience assessment in complex natural systems"

### Figures: S1 to S5
### Table: ST1, ST2

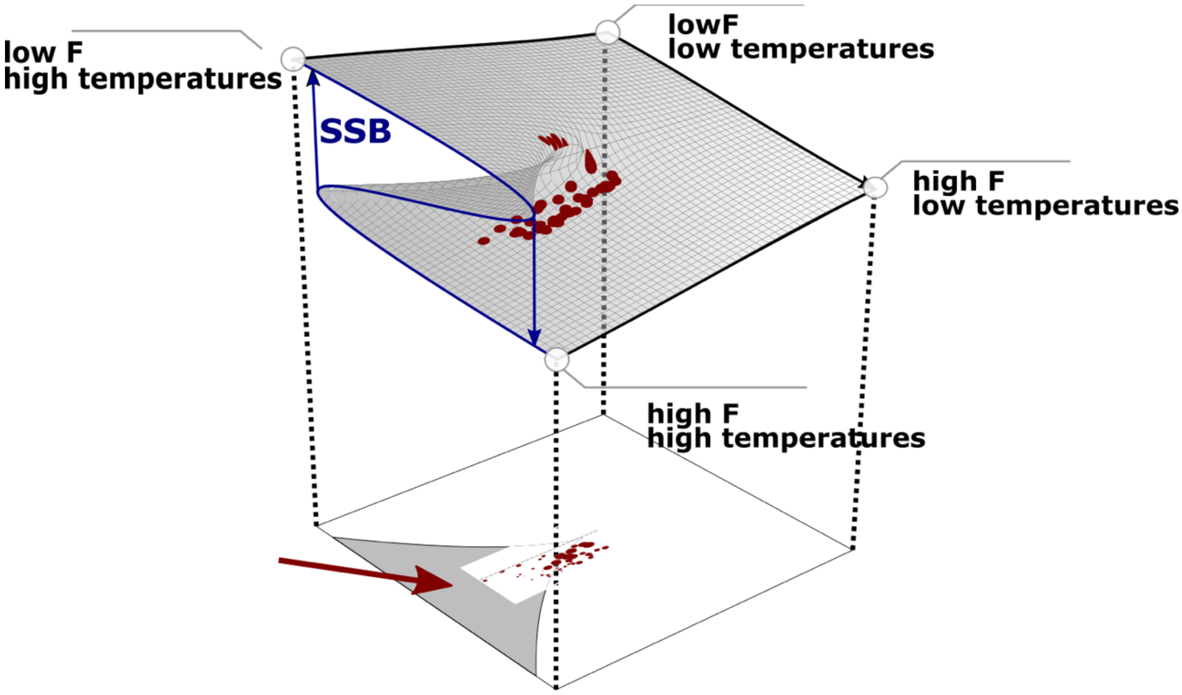


**Figure 1: The stochastic cusp model.** The 3D representation and the 2D projection of the stochastic cusp model. As an example, SSB (Spawning Stock Biomass) was chosen as state variable. Fishing mortality (F) was selected as the splitting factor or the variables controlling whether SSB is high or low. Indeed, when F is low SSB is in the upper shield, while when F is high SSB is in the bottom shield (i.e. low biomass). Temperature was selected as the bifurcation variable controlling the dynamics of SSB. Indeed, the relationship between SSB and F is linear when temperature is high and becomes discontinuous as temperature increase (blue S-shape). The S shape is the typical regime shift curve. Thus the 3D plain, depending on the level of SST and F can represent a linear or a non-linear discontinuous dynamics. The 3D plot can be projected in 2D. The grey area indicated by the arrow is the area below the fold or transition area, where three equilibria are possible two stable and one unstable. The limits of the area correspond to combination levels of the drivers that create a tipping point and favor a regime shift of the state variable. The white area is a linear area, the further away from the transition area the higher the resilience of the state variable. The red dots correspond to the state variable and show how this have changed depending on the two drivers.

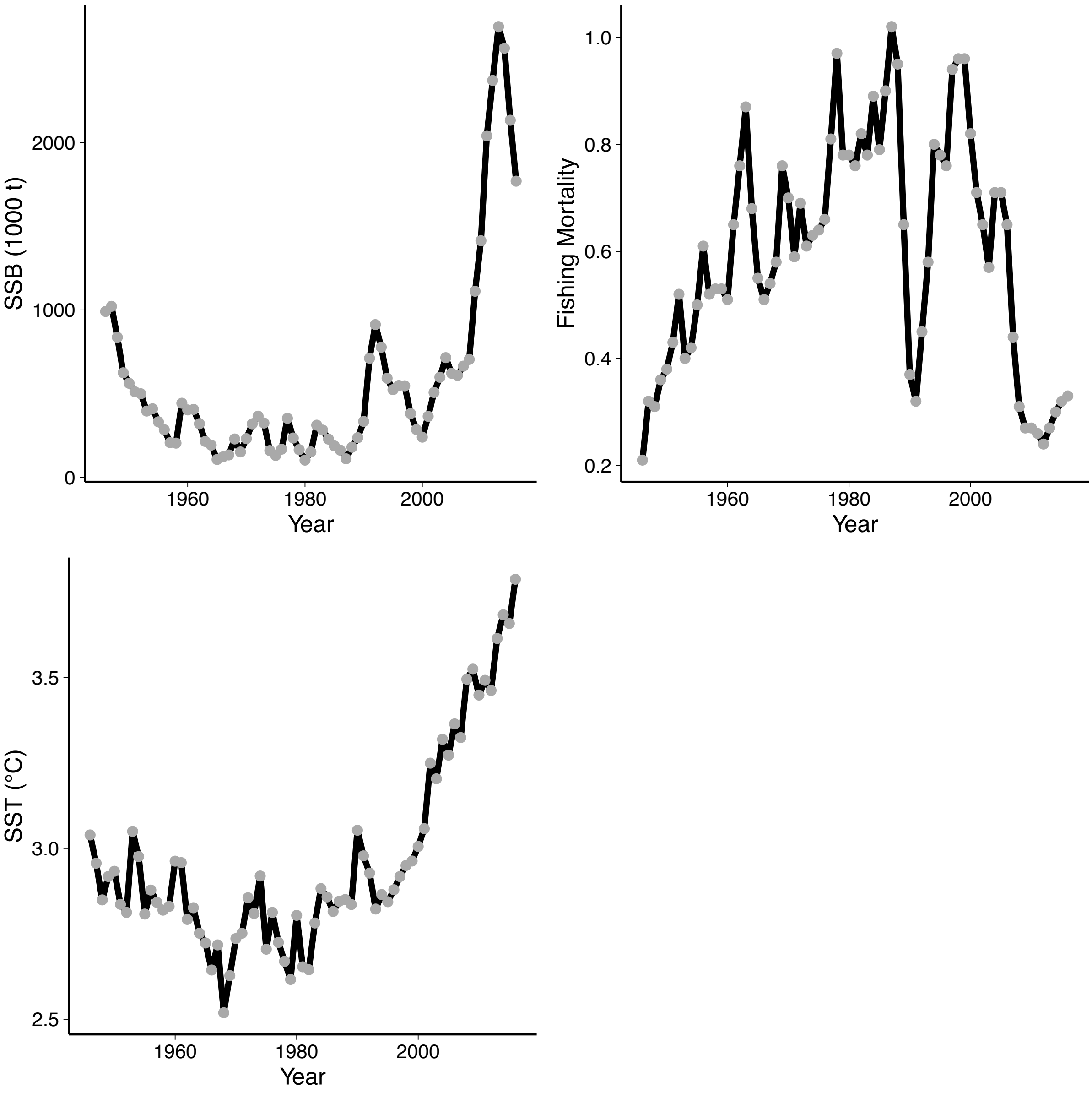


**Figure 2: Time series of the state variable SSB and of the drivers F and SST of North-East Arctic cod.** The time series of Spawning Stock Biomass (SSB) in thousands tonnes, Fishing mortality (F) and Sea Surface Temperature (SST) in °C of North-East Arctic cod from Sguotti et al., 2019.

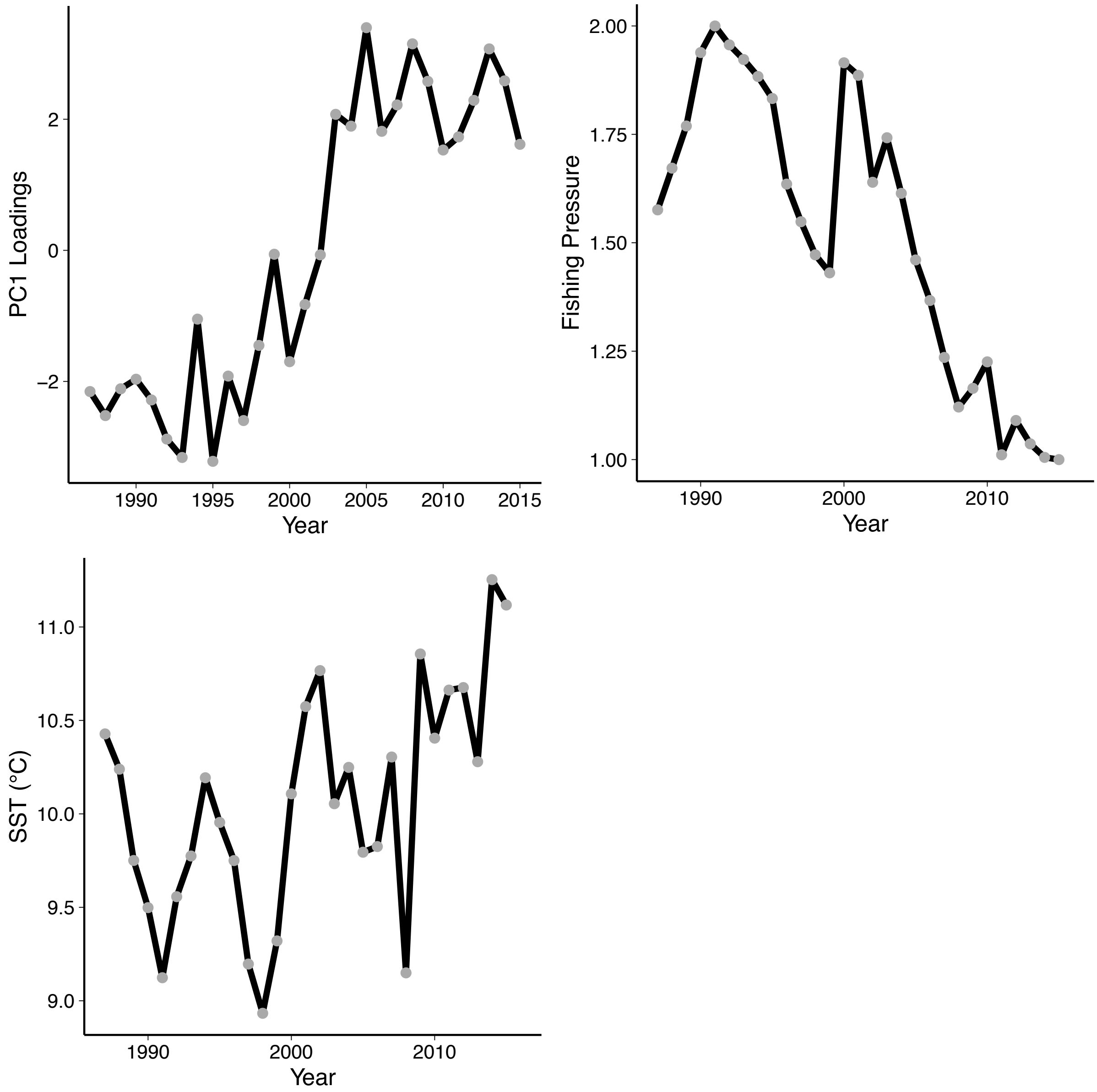


**Figure 3: Variables of the CUSPRA model for the North Sea from Sguotti et al., submitted.** The time series of the loadings of PC1 of the North Sea community, of the fishing pressure and of the Sea Surface Temperature. More info about the data and processing in Sguotti et al., submitted

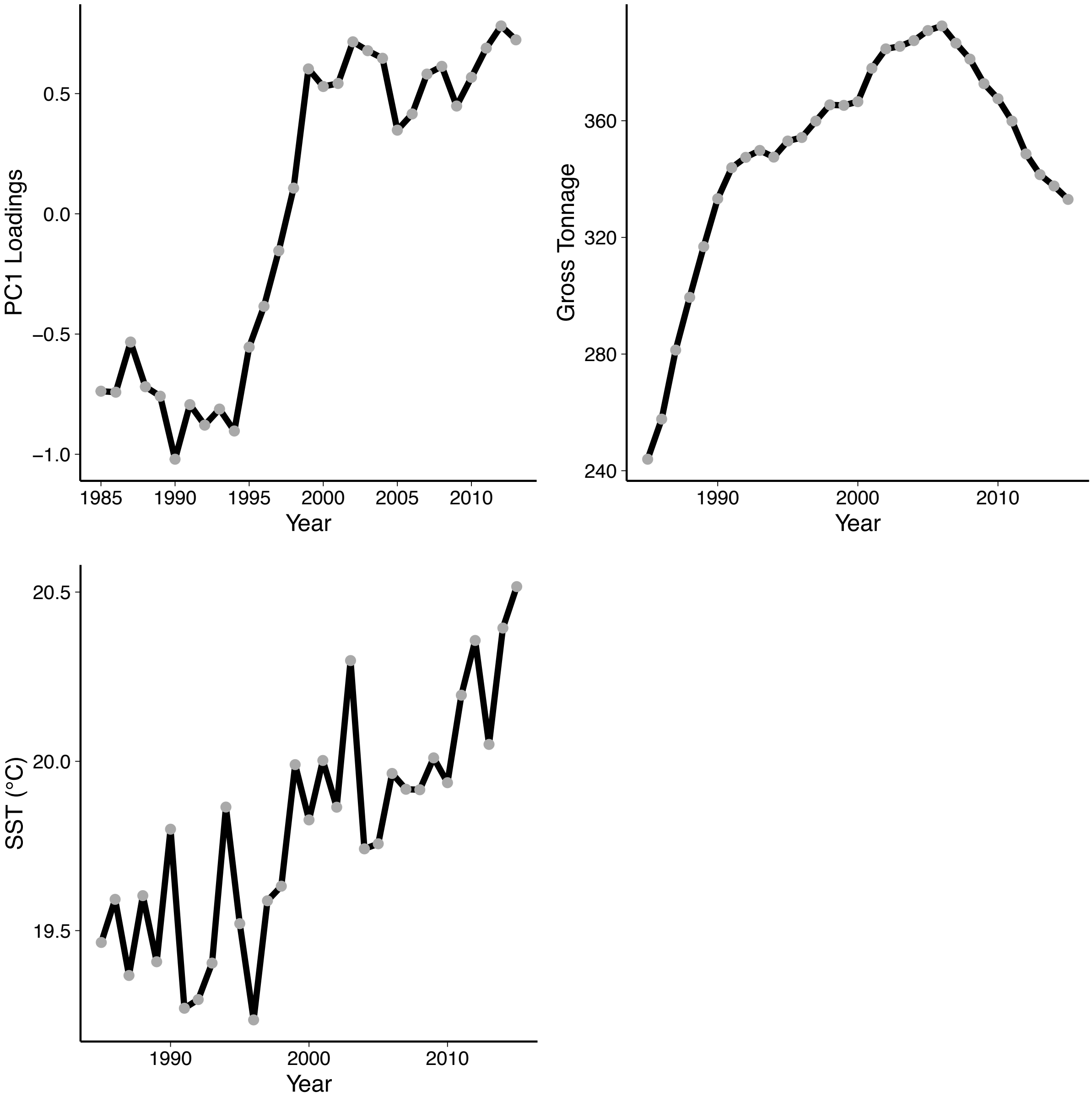


**Figure 4: Variables of the CUSPRA model for the Mediterranean Sea from Vasilakopoulos et al., 2017.** The time series of the PC1 scores of the Mediterranean East Sea community, of the Gross Tonnage used as proxy for fishing capacity and of the Sea Surface Temperature lagged of 2 years. More info about the data and processing in Vasilakopoulos et al., 2017.

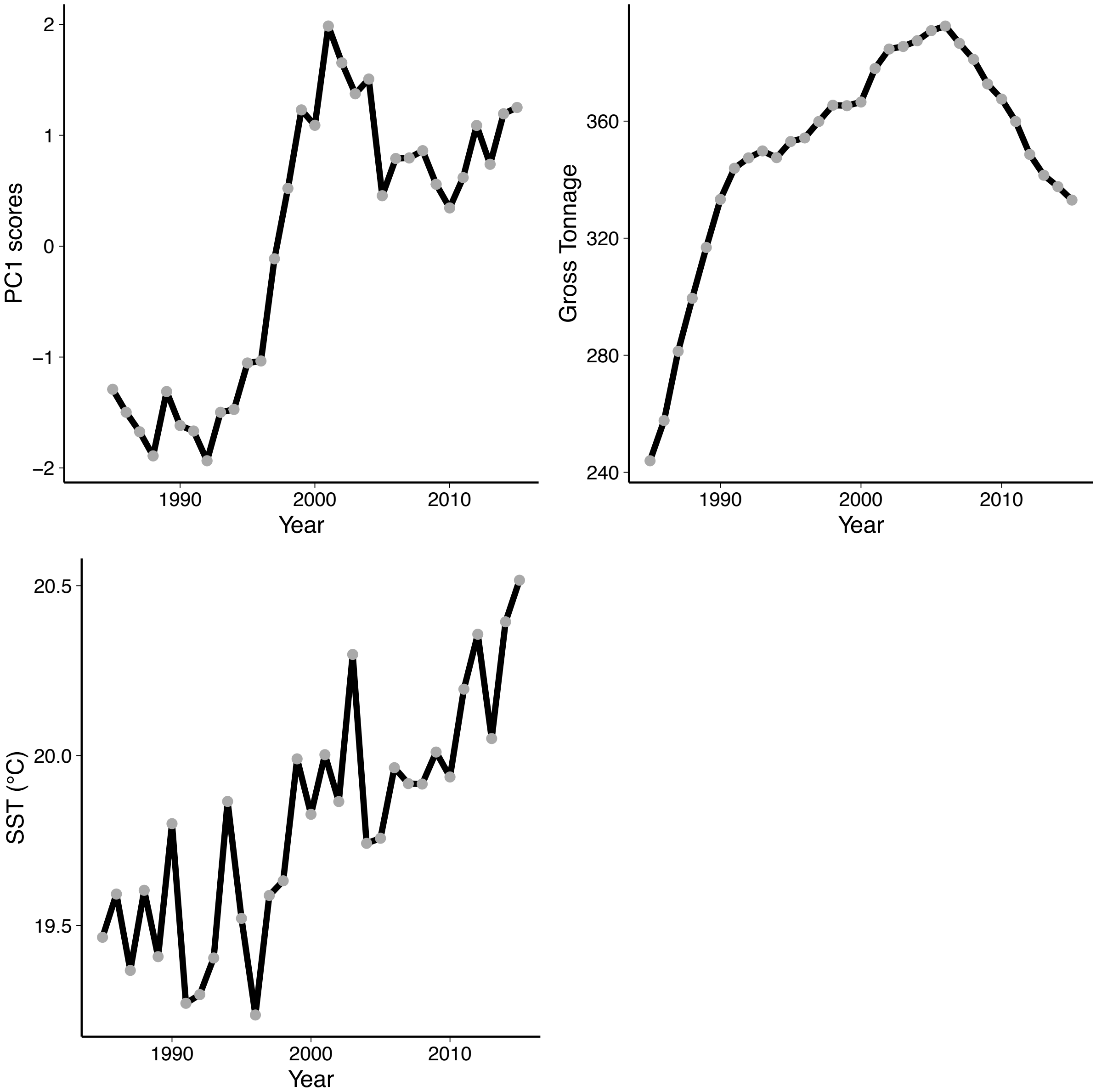


**Figure 5: Variables of the CUSPRA model for the Mediterranean Sea traits from Tsimara et al., 2021.** The time series of the PC1 scores of the trait space of the Mediterranean Sea community, of the Gross Tonnage ad a proxy of fishing capacity and of the Sea Surface Temperature. More info about the data and processing in Tsimara et al., 2021.

**Table1. Results of the CUSPRA method for the 4 examples.** Estimates of the parameters of the stochastic cusp model. $\alpha 1$ and $\alpha 2$ are respectively the intercept and slope of the splitting parameter depending on fishing. $\beta 1$ and $\beta 2$ are respectively the intercept and slope of the bifurcation parameter related to temperature. w1 and w2 are the intercept and the slope of the state variable (in the column name). The significant levels of the estimates are indicated with a star (* $p < 0.05$, ** $p < 0.005$, *** $p < 0.0005$). The $R^2$ are pseudo Cobb $R^2$ computed to compare the model using the cusp formulation or a linear model. Finally, the minimum and maximum levels of RA (resilience from the CUSPRA) are indicated.

| **Parameters** | **North-East Arctic cod** | **North Sea** | **Eastern Med Sea** | **Med Sea Traits** |
|---|---|---|---|---|
| **$\alpha$1** | 2.43 | 6.02** | -8.35** | -8.81** |
| **$\alpha$2** | -9.94 | -3.97** | 0.02** | 0.025** |
| **$\beta$1** | 21.72*** | 12.52* | -74.65* | -13.49 |
| **$\beta$2** | -5.78*** | -1.06 | 3.80** | 0.79 |
| **w1** | -3.18*** | -0.12 | 0.55*** | 0.27 |
| **w2** | 0.002*** | 0.67*** | 2.76*** | 1.16*** |
| **$R^2$ linear** | 0.71 | 0.61 | 0.76 | 0.61 |
| **$R^2$ logistic** | 1 | 1 | 0.82 | 0.71 |
| **$R^2$ cusp** | 0.79 | 0.88 | 0.91 | 0.88 |
| **%point cusp** | >10% | >10% | >10% | >10% |
| **Min RA** | 0 | 0 | 0 | 0 |
| **Max RA** | 1 | 0.99 | 0.95 | 0.94 |

The model output of the stochastic cusp model extended with the CUSPRA is shown for the four examples. The significance of the estimates indicates the extent to which drivers influence the state variable. It is important to check that the estimate of the slope of the state variable (w2) is significant in order for the model to be valid. The pseudo $R^2$ of the cusp model and its alternative linear formulation are calculated and show how well the variables chosen can explain variance in the state variable. The comparison of these two parameters can directly show whether a discontinuous or a continuous dynamic might be better to explain the state variable. Another way to validate the presence of discontinuous dynamics is to look at how many points lie inside the transition area. If more than 10% of the points are inside, the state variable definitely shows a discontinuous dynamic. Finally, the minimum and maximum values of the RA are shown.

**Table 2: Scheme of the case studies used in this paper and the way CUSPRA moves a step forward compared to previous knowledge.**

| Example dataset and main relevant references | Previous knowledge based on application of cusp or IRA | Insight(s) gained by application of CUSPRA | Implications of the insights gained from a management point of view |
|---|---|---|---|
| North-East Arctic cod (Sguotti et al. 2019) | Climate change and fishing induced a positive tipping point with North-East Arctic cod reaching high biomass levels | The stock was resilient but in a low state when temperature was low. The increase of temperature and the decrease of fishing favored a breaking of the resilience and a push of the stock towards another state, now resilient | The stock seems resilient at the moment, but fluctuations in temperature might drive the stock again towards a low resilience status and a new possible shift; thus fishing needs to be limited |
| North Sea ecosystem (Sguotti et al. 2022) | Climate change and fishing pressure induced a regime shift in the North Sea ecosystem in 2003. The ecosystem seems now resilient in a new structure | The ecosystem was not resilient to fishing pressure and climate change from 1985 until 2010. Just now the stock is in a high resilience state and temperature increase will favor its stability | Climate change might have led to irreversible changes in the North Sea, being the ecosystem in a new state and fully resilient. This means that management measures could fail to restore the ecosystem to the previous state, but that caution must be used in management since the system can still switch into something new |
| Eastern Mediterranean fish landings (Vasilakopoulos et al. 2017) | Climate change (sea warming) induced a regime shift in the landed species complex | Increased fishing pressure has pushed the ecosystem into a state of low resilience, but it is the higher temperature that is maintaining the resilience of the system state in low levels | System is currently "trapped" into the area of high instability (low resilience), it could revert to a new state by a drop in temperature, which is highly unlikely. Fisheries management alone cannot push the system state into a new regime. Management should be prepared to manage for a new system state. |
| Mediterranean landings traits (Tsimara et al. 2021) | Climate change (sea warming) induced a regime shift in the configuration of traits landed, as a result of change in species composition. | Increased temperature has pushed the system into a state of low resilience, but it is the reduction in fishing capacity that may be able to move the system state to a new regime | System is currently "trapped" into the area of high instability (low resilience). A reduction of fishing pressure could revert the state; thus, fisheries management is essential for system stability. |
